# ldentify potential feature genes and immune cell infiltration of HIRI based on branched-chain amino acid-related genes by machine learning and experimental validation

**DOI:** 10.1101/2024.12.17.629007

**Authors:** Jiahui Zhao, Min Wu, Shuangjiang Li, Shuang Li, Mingjuan Li, Qiuyun Li, Guangdong Pan, Guoqing Ouyang, Honglai Xu

**Author notes:** These authors have contributed equally to this work as co-first authors (Jiahui Zhao, Min Wu, Shuangjiang Li). These authors have contributed equally to this work as corresponding authors (Guangdong Pan, Guoqing Ouyang, Honglai Xu). **Corresponding author:** Guandong Pan, Department of General Surgery, Liuzhou People’s Hospital Affiliated to Guangxi Medical University, Liuzhou 545006, Guangxi, China Guoqing Ouyang, Department of General Surgery, Liuzhou People ’ s Hospital Affiliated to Guangxi Medical University, Liuzhou 545006, Guangxi, China Honglai Xu, Department of General Surgery, Liuzhou People’s Hospital Affiliated to Guangxi Medical University, Liuzhou 545006, Guangxi, China.

## Abstract

**Background:** Branched-chain amino acid metabolism is involved in the pathogenesis of various liver diseases. In this study, we investigate the potential role of branched-chain amino acid metabolism-related genes in the pathogenesis of hepatic ischemia reperfusion (HIRI).

**Methods:** The gene Expression profiles of HIRI were obtained from the Gene Expression Omnibu database. To determine the differential expression of branched-chain amino acid metabolism-related genes between HIRI and normal tissues. Then, the GO and KEGG analyses were performed, and the protein-protein interaction network was constructed. Next, the random forest and LASSO algorithms were used to screen hub genes, and machine learning techniques were used to build diagnostic models. immunoinfiltration was analyzed in both HIRI patients and controls and the ceRNA network was established. Finally, quantitative real-time PCR was used to verify the expression of hub gene.

**Results:** Based on data set GSE23649, three central DEGs (SLC7A5, SLC1A5, SLC43A2) were determined by the intersection of three machine learning algorithms and used to establish a nomogram that yielded a high predictive performance (area under the curve 0.733−0.922). In the external GSE15480 dataset, AUC value for three key genes is as high as 1.000. Further analysis of nomogram, decision curve and calibration curve also confirme the predictive efficacy of diagnosis. GSEA and GSVA suggest that these three marker genes were involved in multiple pathways associated with HIRI progression. Immunoinfiltration analysis suggest that the proportion of macrophages, neutrophils, aDCs, Treg, and Th1 cells in HIRI group is higher than that in control group, with statistical significance(P<0.05). The ceRNA network demonstrates the complex regulatory relationships among the three hub genes and these mRNA levels were further confirmed in mouse HIRI liver samples.

**Conclusions:** Our study have provided a comprehensive understanding of the association between branched-chain amino acid and HIRI, may provide potential target for HIRI treatment and diagnosis. And provide new insights into the mechanisms of HIRI.

**Graphical Abstract:** 

## Introduction

During liver resection, ischemia-reperfusion (I/R) is a necessary procedure to minimize intraoperative bleeding. However, this process invariably causes liver damage and can lead to severe postoperative complications, including liver failure1. The widespread use of liver transplantation and hepatectomy has heightened interest in strategies to prevent and mitigate hepatic ischemia-reperfusion injury (HIRI). The mechanisms underlying HIRI involve both direct cellular damage due to ischemia and delayed injury caused by the activation of inflammatory pathways2. HIRI progresses in two distinct phases: ischemia and reperfusion. In the ischemic phase, reduced blood flow and oxygen deprivation in liver tissues lead to the accumulation of reactive oxygen species (ROS). This increase in ROS triggers oxidative stress, contributing to endothelial dysfunction, DNA damage, and localized inflammation. These inflammatory cascades and oxidative stress can provoke a cytokine storm, leading to cell death and tissue damage3. In the reperfusion phase, the restoration of blood flow to ischemic tissues results in the release of phospholipase A2, TNF-α, IL-1β, IFN-γ, and angiotensin II, which activate NADPH oxidase. Phospholipase A2 promotes the production of platelet-activating factor, increasing thromboxane and leukotrienes, which exacerbate local inflammation4–6. Additionally, angiotensin II stimulates its receptors, upregulating NADPH oxidase expression and contributing to ischemia-reperfusion injury through the angiotensin-converting enzyme pathway7,8. Despite extensive basic and clinical research, the precise mechanisms underlying ischemia-reperfusion injury remain to be further clarified.

Branched-chain amino acids (BCAAs), comprising valine, leucine, and isoleucine, are essential for human nutrition. BCAAs are known to promote skeletal muscle protein synthesis, enhance protein metabolism, and inhibit protein degradation9. Previous study have demonstrated that BCAAs stimulate mTOR mRNA translation, thereby regulating protein synthesis at the molecular level10. BCAAs also mitigate hepatic steatosis and liver injury associated with nonalcoholic steatohepatitis by inhibiting the expression of the FAS gene(1)11. Furthermore, BCAAs can block the Akt2-INSIG2a signaling pathway through the mTOR pathway, reducing intrahepatic adipogenesis and redistributing lipids in the liver, muscles, and kidneys12. Recent research also indicate that fructose ingestion induces the hepatic transcription factor ChREBP-β, which activates BCKDK transcription, thereby linking fructose metabolism with BCAA metabolism13. Although significant progress has been made in understanding the relationship between BCAA metabolic regulation and disease, the potential role of BCAAs in HIRI remains unclear, further research is needed to explore the potential role of BCAA as a therapeutic target for HIRI.

In recent years, bioinformatics and machine learning strategies have advanced rapidly, enabling a deeper exploration of underlying mechanisms and potential biomarkers. In this study, we analyze the differential expression and immunological characteristics of BCAA-related genes(BRGs) in 33 cases of HIRI and 33 control cases. Machine learning algorithms were employed to identify key genes for diagnostic prediction. We validated the predictive model using a nomogram, decision curve analysis (DCA), calibration curve, and receiver operating characteristic (ROC) curve. Additionally, we explored the relationship between hub BRGs and immune infiltration and constructed a ceRNA network.

## Materials and methods

### Data collection and processing

The GSE23649 dataset was retrieved from the NCBI Gene Expression Omnibus(GEO; https://www. ncbi.nlm.nih.gov/geo/) using the keywords “Hepatic ischemia reperfusion.” This dataset, used as the training set, comprises 33 hepatic ischemia-reperfusion samples (9 living donor samples, 8 cardiac death donor samples, 8 brain death donor samples with early allograft dysfunction, and 8 brain death donor samples without early allograft dysfunction) and 33 control samples. The GSE15480 dataset served as the validation set, consisting of 6 control samples and 6 HIRI samples. The flowchart of this study is shown in Figure 1.

**Figure 1.**
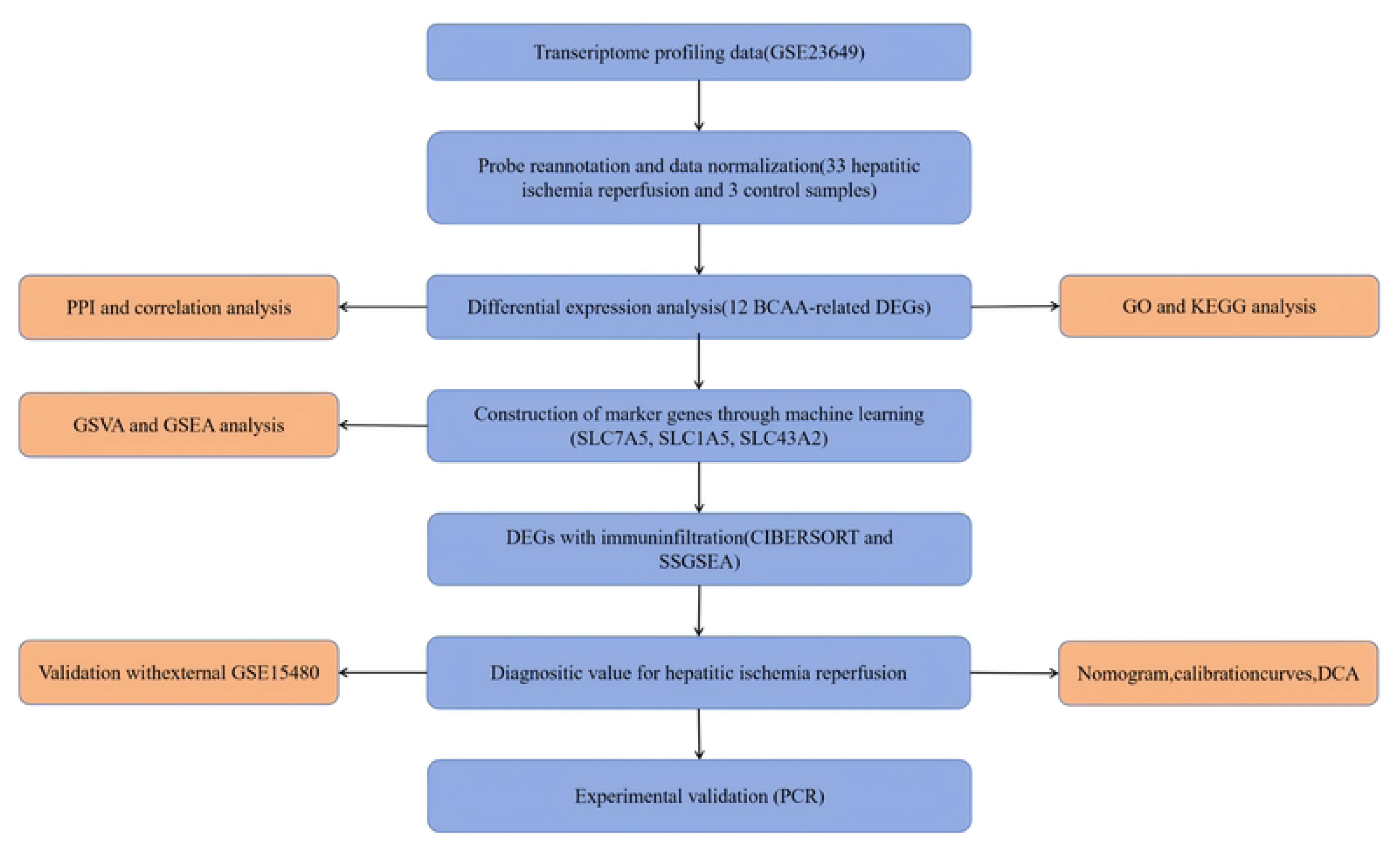
Flowchart of the present study.

### Differential gene expression analysis of DEGs

Differentially expressed genes (DEGs) related to branched-chain amino acids (BCAA) between the hepatic ischemia-reperfusion and control groups were identified using the Wilcoxon signed-rank test. To enhance the visualization of these DEGs, the “ggplot2” and “pheatmap” R packages were employed to generate boxplots and heatmaps. The “VennDiagram” R package was used to determine the intersection of BRGs, which were defined as the BRGs for subsequent analyses. Statistical significance was set at p < 0.05.

### Correlation analysis and PPI network construction

Landscape plots of the 23 chromosomes and a heatmap of the 13 DE-BRGs were generated using the “RCircos” and “heatmap” R packages, respectively. Pearson ’ s correlation analysis was performed to create Circos plots of DE-BRGs using the “ circlize ” package. A protein-protein interaction (PPI) network for the 13 DE-BRGs was constructed using the STRING database(https://string-db.org/), with a medium confidence threshold of 0.4.

### Pathway and functional enrichment analysis

The Kyoto Encyclopedia of Genes and Genomes (KEGG) database was utilized as a comprehensive resource for the systematic analysis of gene functions14. Gene ontology (GO) analysis identified the biological processes (BP), cellular components (CC), and molecular functions (MF) associated with the 13 DE-BRGs. The “clusterProfiler” R package was used for both GO and KEGG pathway analyses, with visualizations generated using the “enrichplot” R package. The significance threshold for enrichment was set at p < 0.05.

### Construction of DEGs diagnostic model

The GSE23649 dataset was used as the training set, while the GSE15480 dataset was used as the validation set for the machine learning model. Three algorithms—random forest (RF), support vector machine-recursive feature elimination (SVM-RFE), and least absolute shrinkage and selection operator (LASSO) regression—were employed to identify the most robust hub genes for diagnosing hepatic ischemia-reperfusion injury (HIRI). LASSO regression, implemented using the “glmnet” R package, reduced data dimensionality while preserving valuable variables15,16. SVM-RFE is a method for detecting optimal variables in machine learning by pruning support vectors. If there are many features, halve the features for each round17. The “e1071” R package was used to construct the SVM-RFE models for screening the best variable genes. The RF model is an ensemble machine learning method for determining the optimal number of variables using various independent decision trees18. The result can be obtained using the “randomForest” R package. The intersection of the RF, LASSO algorithm, and SVM-RFE results was taken to yielded the most significant hub genes.

### ROC evaluation and nomogram construction

To assess the predictive accuracy of each hub gene, receiver operating characteristic (ROC) curves were generated, and the area under the curve (AUC) along with 95% confidence intervals (CI) were calculated using the “pROC” R package. The external GSE15480 dataset was used to validate the diagnostic model. A nomogram was constructed using the “rms” R package, where each hub gene was assigned a score. The total score predicted the likelihood of HIRI occurrence. Clinical impact curves and decision curves were also generated to evaluate the model’s clinical efficacy.

### Gene set enrichment analysis and gene set variation analysis

To explore the potential functions of the hub genes, we conducted gene set enrichment analysis (GSEA) using the KEGG gene set (c2.cp.kegg.symbols.gmt) and GO gene set (c5.go.symbols.gmt) from the Molecular Signatures Database. The “clusterProfiler” R package was used for GSEA, while gene set variation analysis (GSVA) was performed using the “GSVA” R package to identify differentially enriched pathways between high and low expression subtypes based on hub gene expression. Differential expression pathways were further identified using the “limma” R package.

### Analysis of immune cell infiltration

The CIBERSORT algorithm was applied to convert normalized gene expression matrices into immune cell components19. We analyzed the proportion of 22 immune cells in the GSE23649 dataset, with a p-value < 0.05 defining accurate immune cell fractions. Correlation heatmaps of immune cells in HIRI pathogenesis were generated using the “corrplot” R package, and boxplots were used to compare the expression levels of different immune cells. Spearman correlation analysis was conducted to determine associations between hub genes and immune infiltrating cells using the “GSVA” R package.

### ceRNA network construction

We use miRanda (http://www.microrna.org/), miRDB (http://www.mirdb.org/) and TargetScan (https://www.targetscan.org/ The vert 80/) database to predict miRNA-mRNA interactions between these three hub genes. Sponge scan (http://spongescan.rc.ufl.edu/) was used to predict the target of the direct interaction between miRNA and lncRNA. Then, the mRNA-miRNA-lncRNA ceRNA network was established and visualized by Cytoscape software (version 3.9.0).

### HIRI mouse model establishment and histological procedure

C57BL/6J mice were obtained from Hunan Slack Jingda Laboratory Animal Co., LTD. Mice had unrestricted access to sterile water and food. In 8-week-old mice, hepatic ischemia was induced by clamping the hepatic artery and portal vein for 30 minutes, followed by reperfusion. Mice were euthanized after 6 hours of reperfusion, and serum and liver samples were collected. Serum ALT and AST levels determined by a fully automated biochemical immunoassay system (cobas® 8000, Germany). Fresh liver tissue was immersed in a 4% paraformaldehyde solution and used for paraffin embedding.4 mm thick sections were obtained using a microtome and subsequently stained with H&E according to the manufacturer’s protocol. All animal experiments were approved by the Animal Care and Use Committee of Liuzhou People’s Hospital Affiliated to Guangxi Medical University.

### RNA isolation and real-time quantitative PCR

RNA was extracted from HIRI mouse liver tissues using TRIzol reagent (15596026CN, thermofisher, Guangzhou, China). RNA was reverse transcribed using the ReverAid™ Master Mix with Dnase I-LBID(M16325,thermgfisher, Guangzhou).

Real-time quantitative PCR (qRT-PCR) was performed using the lightCysler @96 automatic fluorescence quantitative PCR instrument (Roche Diagnostic Products Co.Ltd., Shanghai, China) and PowerUp ™ SYBR ™ Green (A25742, thermofisher, Guangzhou). Primer sequences are listed in Supplementary Table S1.

### Statistical analysis

Statistical and data analyses were performed using R software (version 4.2.1).

Continuous data are expressed as mean ± standard deviation. For comparisons between two groups, the Student’s t-test was applied to normally distributed variables, while the Wilcoxon rank-sum test was used for non-normally distributed variables. A p-value less than 0.05 was considered statistically significant. ns, P > 0.05; *, P < 0.05; **, P < 0.01; ***, P < 0.001.

## Results

### Differential expression genes identification in HIRI

With the “limma” R package, a total of 5110 Differentially expressed genes (DEGs) were identified based on the HIRI dataset GSE23649, of which 2,701 were downregulated and 2,409 were upregulated (Figure 2A). The gene expression patterns of 5110 DEGs were shown in the heatmap (Figure 2B). The intersection of 50110 DEGs and 33 genes related to BCAA metabolism revealed 13 BCAA-related DEGs (DE-BRGs) (ABAT, ACAD8, ACADSB, AUH, BCKDHB, DLD, ECHS1, IVD, MCCC1, SLC1A5, SLC3A2, SLC43A2, SLC7A5) with significant differences between HIRI and control group (Figure 2C). The chromosomal locations of the 13 DE-BRGs are shown in Figure 2D. We found that SLC7A5, SLC1A5, SLC3A2, and SLC43A2 were up-regulated in HIRI, while DLD, AUH, MCCC1, ACAD8, IVD, ABAT, ACADSB, ECHS1, and BCKDHB were down-regulated (Figures 2E, F). The correlation among the 13 DE-BRGs is shown in Figure 2G-H. MCCC1 was positively correlated with ABAT, ACAD8 and BCKDHB, negatively correlated with SLC1A5 and SLC3A2. To investigate the potential relationship between these 13 DE-BRGs, we performed PPI analysis as shown in Figure 2I.

**Figure 2.**
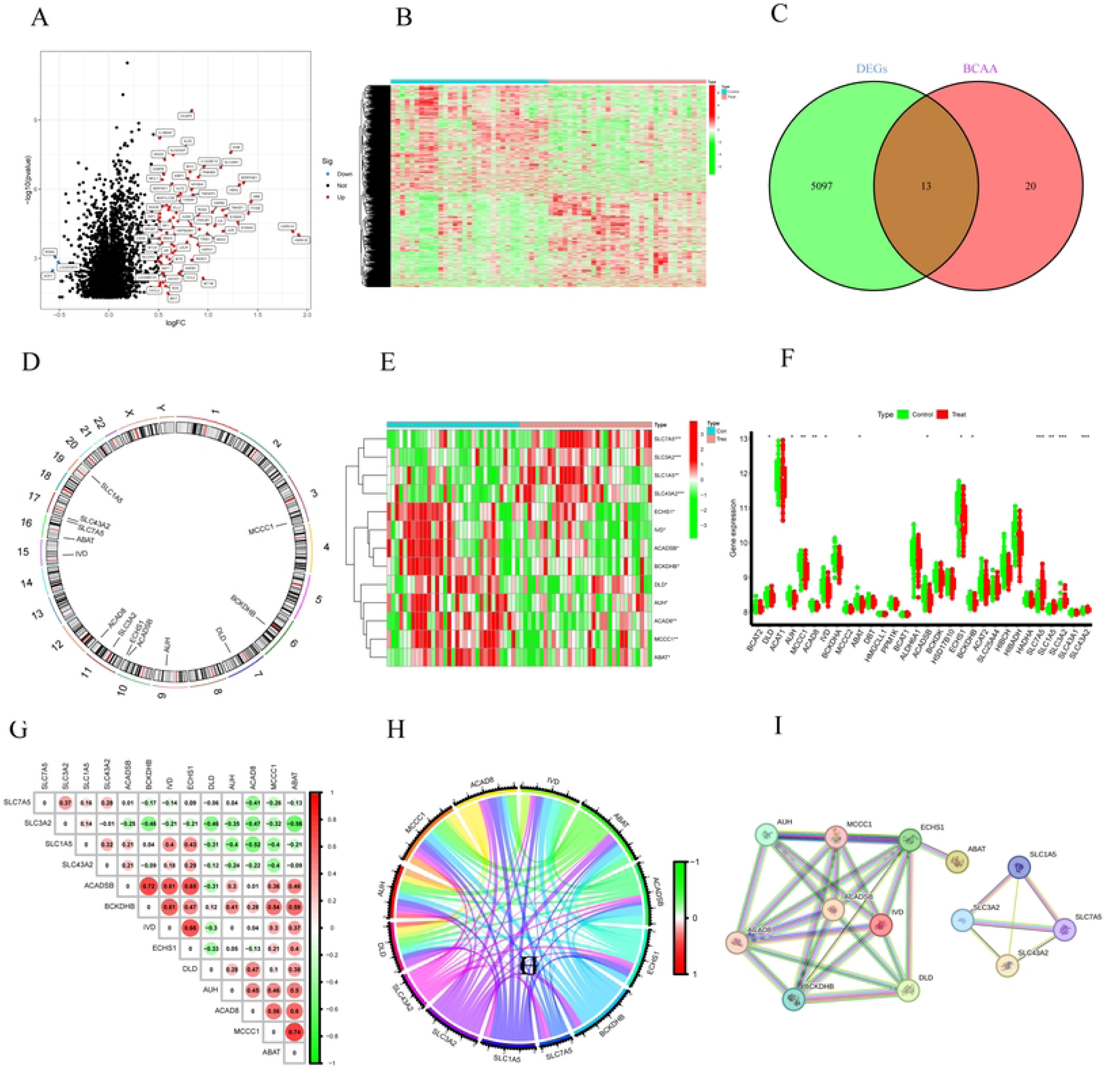
Identification of BCAA-related differentially expressed genes in HIRI. (A) DEGs are shown by volcano plots, with blue denoting the downregulated genes, red denoting the upregulated genes. (B) The gene expression patterns of DEGs were shown in the heatmap. (C) Venn diagram showed the intersection of genes between DEGs and BCAA-related genes. (D) The locations of the 13 DE-BRGs on 23 chromosomes. (E) The expression patterns of 13 DE-BRGs were shown in the heatmap. (F) Boxplot showed the differential expression of DE-BRGs between HIRI and control samples. (G) Correlation analysis of 13 DE-BRGs. Red and green colors represent positive and negative correlations respectively. (H)Correlation analysis of 13 DE-BRGs in the circle graph. Red and green colors represent positive and negative correlations respectively. (I)Gene relationship network diagram of 13 DE-BRGs. p values were showed as: *, p < 0.05; **, p < 0.01; ***, p < 0.001. DEGs, differential expression genes; DE-BRGs, differential expression of branched-chain amino acid metabolism-related genes.

### Functional enrichment analysis of DE-BRGs in HIRI

To gain deeper insights into the signaling pathways involved in HIRI and DE-BRGs, we conducted pathway and functional enrichment analyses using the “ClusterProfiler” R package to elucidate the associated biological functions and pathways. The Gene Ontology (GO) enrichment analysis revealed that the most significant biological processes (BP) included the response to branched-chain amino acid catabolic process, branched-chain amino acid metabolic process, amino acid catabolic process, and organic acid catabolic process. Cellular component (CC) analysis highlighted significant involvement in the mitochondrial matrix, basal plasma membrane, basal part of the cell. Molecular function (MF) analysis identified strong associations with neutral L-amino acid transmembrane transporter activity, L-amino acid transmembrane transporter activity, amino acid transmembrane transporter activity, and carboxylic acid transmembrane transporter activity (Figure 3A). Additionally, KEGG pathway analysis indicated that the 13 DE-BRG were notably enriched in pathways such as valine, leucine, and isoleucine degradation, propanoate metabolism, lipoic acid metabolism, and 2-oxocarboxylic acid metabolism (Figure 3B). These findings align with the results of the GO enrichment analysis.

**Figure 3.**
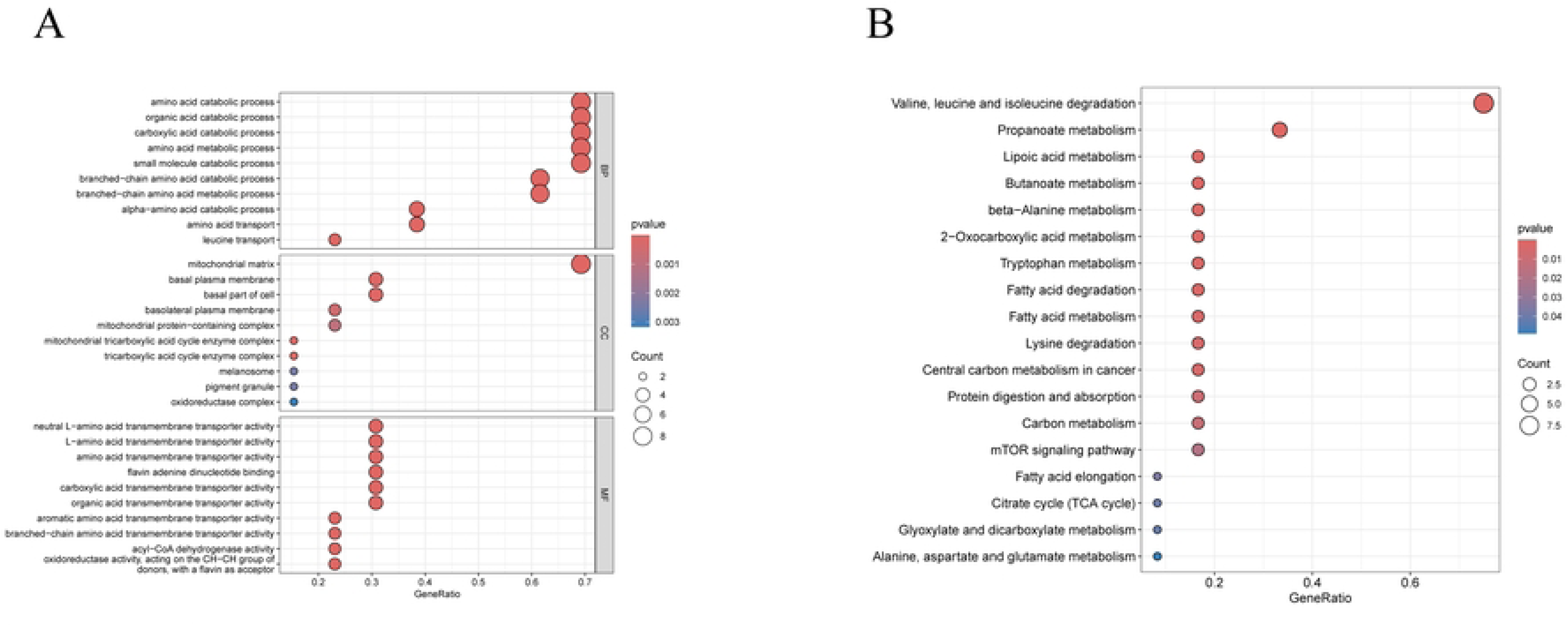
Functional analyses for the DE-BRGs. (A) Bubble diagrams of the GO enrichment analysis of 13 DE-BRGs. (B) Bubble diagrams of the KEGG enrichment analysis of 13 DE-BRGs. DE-BRGs, differential expression of branched-chain amino acid metabolism-related genes. GO, Gene Ontology; BP, Biological process; CC, Cellular component; MF, Molecular function; KEGG, Kyoto Encyclopedia of Genes and Genomes.

### Construction of the diagnostic hub genes for HIRI

To identify the hub genes with the highest diagnostic value, we employed LASSO regression and two validated machine learning models—support vector machine-recursive feature elimination (SVM-RFE) and random forest (RF)—to filter the 13 DE-BRGs for their predictive potential in diagnosing HIRI. LASSO logistic regression identified 8 key genes from the DE-BRGs (Figure 4A-B). The RF model selected all 8 DE-BRGs (Figure 4C-D). The SVM-RFE model achieved the smallest error and highest accuracy (minimal error = 0.391, maximal accuracy = 0.609) when selecting 5 features: SLC7A5, SLC43A2, BCKDHB, SLC1A5, IVD (Figure 4E-F). The intersection of genes identified by the RF, LASSO, and SVM-RFE models revealed three hub genes—SLC7A5, SLC1A5, and SLC43A2—that were considered to possess the highest diagnostic value (Figure 4G).

**Figure 4.**
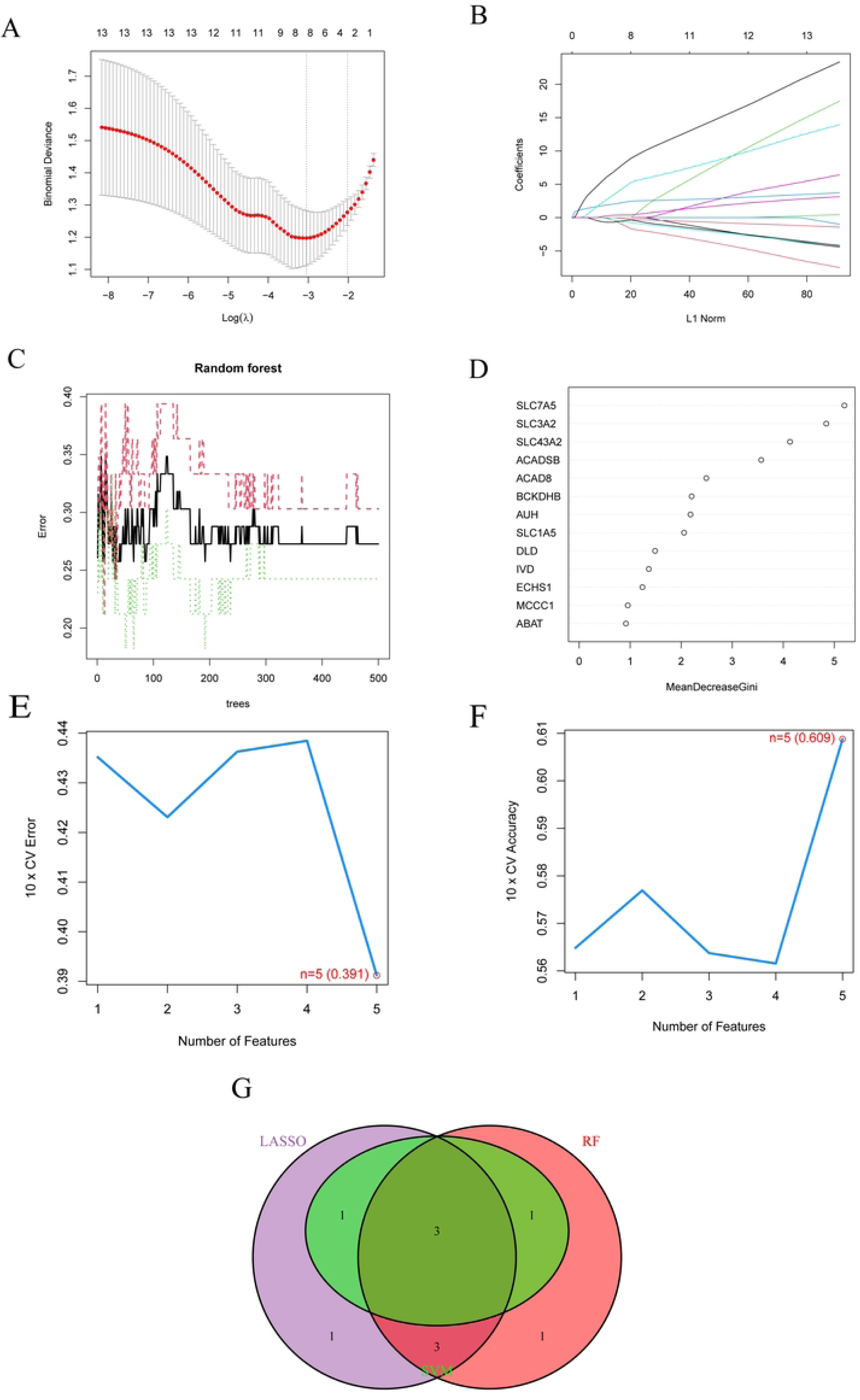
Candidate biomarker identification for HIRI via machine learning algorithms. (A) LASSO regression coefficient analysis. The solid vertical lines represent the partial likelihood deviance SE. The dotted vertical line is drawn at the optimal lambda. (B) LASSO regression cross validation graph. Each curve corresponds to one gene. (C) Relationship between the number of random forest trees and error rates. The red line represents the error of the HIRI group, the green line represents the error of the Control group, and the black line represents the total sample error. (D) The rank of genes in accordance with their relative importance. (E) The error of the feature selection for the SVM-RFE algorithm, with the lowest cross-validation error is found in 4 gens and the values is 39.1%. (F) The accuracy of the feature selection for the SVM-RFE algorithm. The peak of the curve is achieved at 4 genes with an accuracy of 60.9%. (G) The Venn diagram shows the overlap of marker genes between LASSO, random forest, and SVM-RFE algorithms.

### Evaluation of the predictive value and nomogram construction

ROC curve analysis demonstrated robust predictive performance for each of the hub genes, with area under the curve (AUC) values as follows: SLC7A5 (AUC: 0.791), SLC1A5 (AUC: 0.729), and SLC43A2 (AUC: 0.761)(Figure 5A). The overall model achieved an AUC of 0.834 (95% CI: 0.733 − 0.922) (Figure 5B). To further assess the predictive efficiency of these three hub genes, we constructed a nomogram model for HIRI patients using the “rms” R package (Figure 5C).In this nomogram, the relative expression of each gene corresponds to a score, and the total score—calculated by summing the scores of all genes—predicts the risk of HIRI. The calibration curve indicates the relative relationship between the predicted and actual probabilities, with closer solid and dashed lines indicating higher model accuracy. The predicted curve matches the standard curve well, indicating that the nomogram model for HIRI is accurate (Figure 5D). DCA indicated that the “ Model” curve was higher than the gray line, suggesting a high level of accuracy and offered a better clinical benefit(Figure 5E). The clinical impact curve further confirmed this result (Figure 5F).

**Figure 5.**
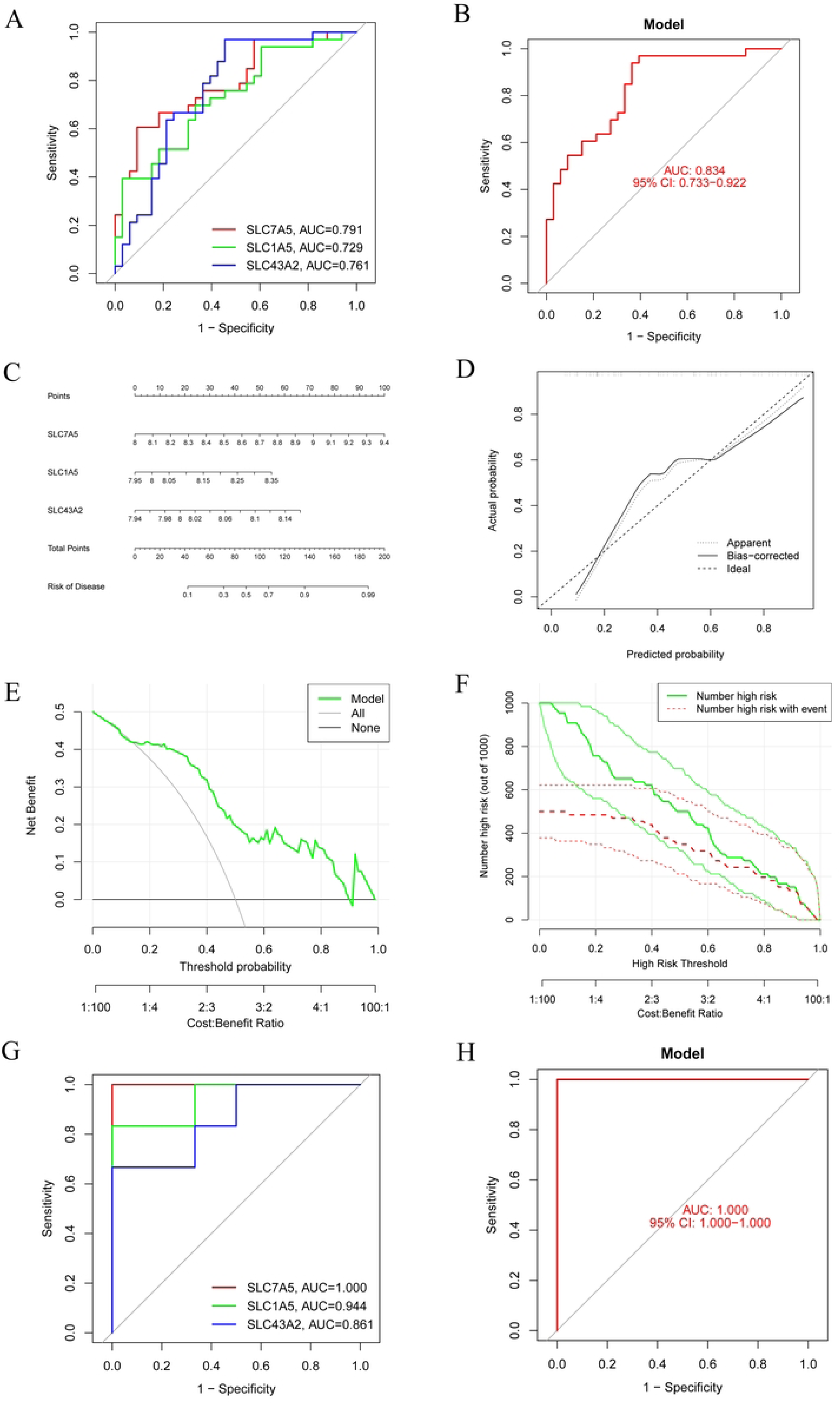
Validation of biomarker gene expression (A) The ROC results for the 3 marker genes. The AUC value of SLC7A5, SLC1A5 and SLC43A2 was 0.791, 0.729, 0.761, espectively. (B) The ROC curves were evaluated comprehensively by GSE23649. The AUC value was 0.834(95% CI: 0.733−0.922). (C) Nomogram graph of the 3 marker genes. (D) Calibration curve displaying the diagnostic ability of the nomogram model. (E) DCA illustrating the predictive efficiency of Nomogram models. (F) The clinical impact curve showed a higher diagnostic ability of the nomogram model. (G) The ROC results of 3 marker genes in validation set. The AUC value of SLC7A5, SLC1A5 and SLC43A2 was1.000, 0.944, 0.861, respectively. (H) The ROC curves of the three hub genes were evaluated comprehensively by the validation set. The AUC value was 1.000(95% CI: 1.000−1.000). AUC, area under curve; ROC, receiver operating characteristic; DCA, Decision curve analysis.

To further validate the model’s ability to identify HIRI patients, we tested its performance using the GSE15480 dataset. The result showed that SLC7A5, SLC1A5, and SLC43A2 in HIRI group was upregulated compared with control group (Figure 5G). The AUC value of ROC for SLC7A5, SLC1A5, and SLC43A2 were 1.0, 0.944, 0.861, respectively. Additionally, the AUC value of model was 1.0, confirming the model’s strong predictive accuracy (Figure 5H). These findings suggest that the three identified hub genes—SLC7A5, SLC1A5, and SLC43A2—may serve as reliable diagnostic markers for HIRI.

### Profile of GSEA and GSVA

To elucidate the key signaling pathways associated with hub genes in hepatic ischemia-reperfusion injury (HIRI), we performed single-gene Gene Set Enrichment Analysis (GSEA) using KEGG and GO pathway datasets. The GSEA of KEGG pathways revealed that high expression of SLC43A2, SLC1A5, and SLC7A5 was significantly associated with antigen processing and presentation. Additionally, SLC43A2 and SLC7A5 were linked to cytokine-cytokine receptor interaction, the MAPK signaling pathway, and the NOD-like receptor signaling pathway (Figure 6A-B). Both SLC1A5 and SLC7A5 were associated with systemic lupus erythematosus. In contrast, high expression of SLC1A5 was connected to the mTOR signaling pathway, N-Glycan biosynthesis, the proteasome, ECM-receptor interactions, and systemic lupus erythematosus (Figure 6C). SLC7A5 was also notably associated with complement and coagulation cascades. The GO enrichment results from GSEA are presented in Figures 6D-F.

**Figure 6.**
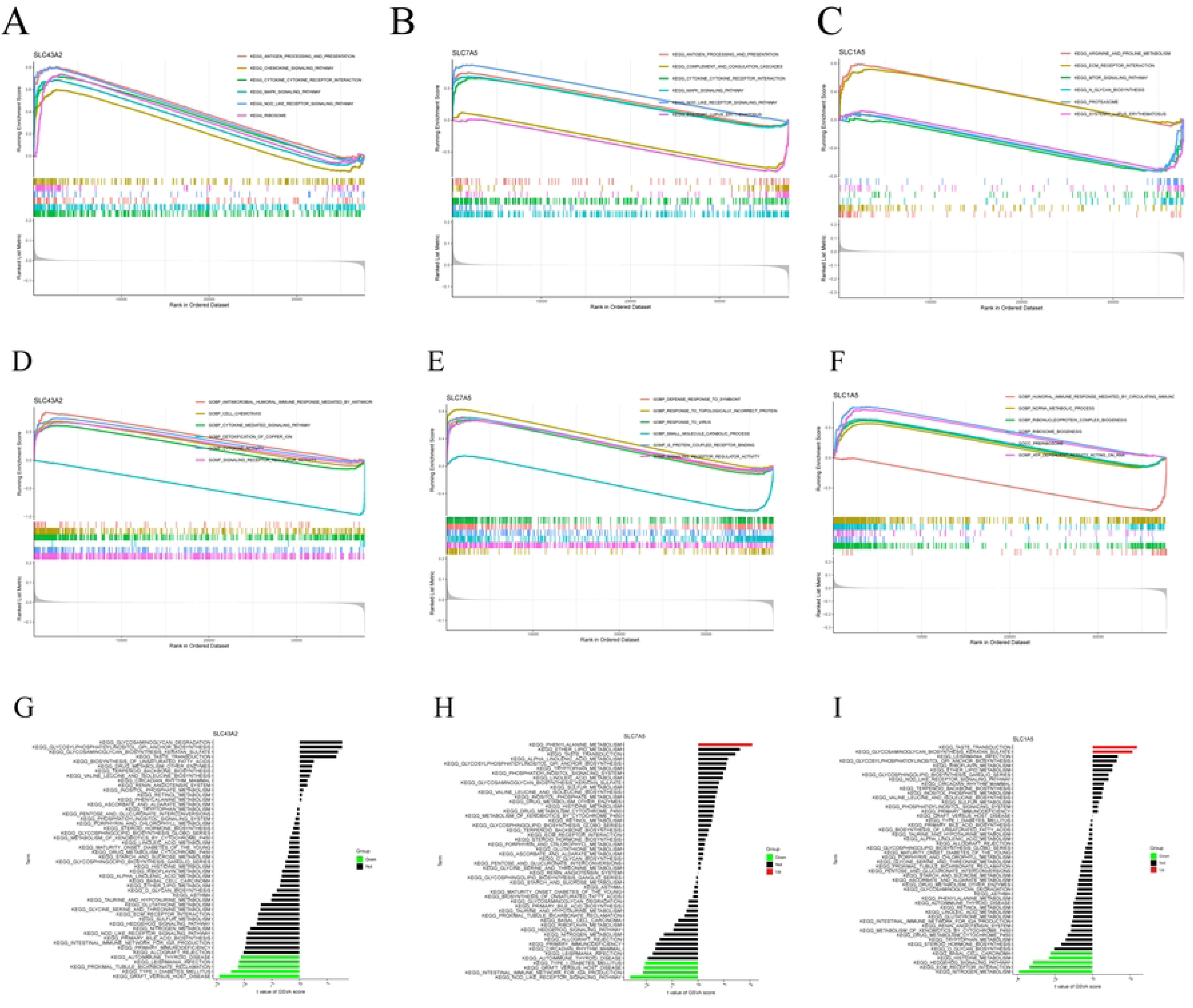
GSEA and GSVA analysis of three marker genes. The KEGG and GO pathway enrichment analysis of (A, D) SLC43A2, (B, E) SLC7A5 and (C, F) SLC1A5 were carried out by GSEA enrichment method, and the two items with the highest and lowest enrichment scores are visualized according to the arrangement of enrichment scores. The KEGG pathway enrichment analysis of (G) SLC43A2, (H) SLC7A5 and (I) SLC1A5 was carried out by GSVA enrichment method, and the top 50 are visualized according to the enrichment score.

We then employed GSVA to identify differentially active pathways between low- and high-expression subtypes based on the expression levels of the three hub genes. The results indicated that the expression of both SLC7A5 and SLC43A2 were associated with Type I diabetes mellitus and graft versus host diseases. The low expression of SLC1A5 is implicated in pathways such as nitrogen metabolism, ECM receptor interaction, hedgehog signaling pathway, and basal cell carcinoma. Downregulated of SLC7A5is associated with graft versus host disease, intestinal immune network for IGA production, and NOD-like receptor signaling pathway. Whereas, decreased expression of SLC43A2 were correlated with autoimmune thyroid disease, leishmania infection (Figure 6G, I). High expression of SLC7A5 was significantly involved in the phenylalanine metabolism (Figure 6H). Additionally, high expression of SLC1A5 was correlated with taste transduction and glycosaminoglycan biosynthesis keratan sulfate (Figure 6I).

### Immunoinfiltration analysis

To explore whether BRGs could influence HIRI progression by modulating immune infiltration, we employed single-sample GSEA (ssGSEA) to investigate the correlation between these hub genes and immune cells and functions infiltration in HIRI. The heatmap show the distribute of immune cells in control and HIRI group (Figure7A). As for immune cells, the ssGSEA algorithm revealed that aDCs, Macrophages, Neutrophils, Th1 cells, and Treg were upregulated in HIRI group. In term of immune functions, the ssGSEA scores of APC co-inhibition, CCR, inflammation promoting, MHC class I, T cell co-inhibition, Type I IFN response were higher in HIRI group than control group, while cytolytic activity and Type II IFN response were higher in control group (Figure7C).

Figure 7B and Supplementary Table S2 showed the correlation between 3 hub genes and immune infiltration. The result showed that SLC1A5 was significantly related to T helper cells, T cell co-stimulation, APC co-inhibition. SLC43A2 associated with aDCs, CCR, MHC class I, Type I IFN response, Parainflammation, APC co-stimulation. In addition, SLC7A5 significantly correlated with aDCs, CCR, Neutrophils, Parainflammation, Treg. Figure 7D shows the correlation between immune cells. These findings suggest that the three hub genes are closely linked to the immune infiltration microenvironment in HIRI.

**Figure 7.**
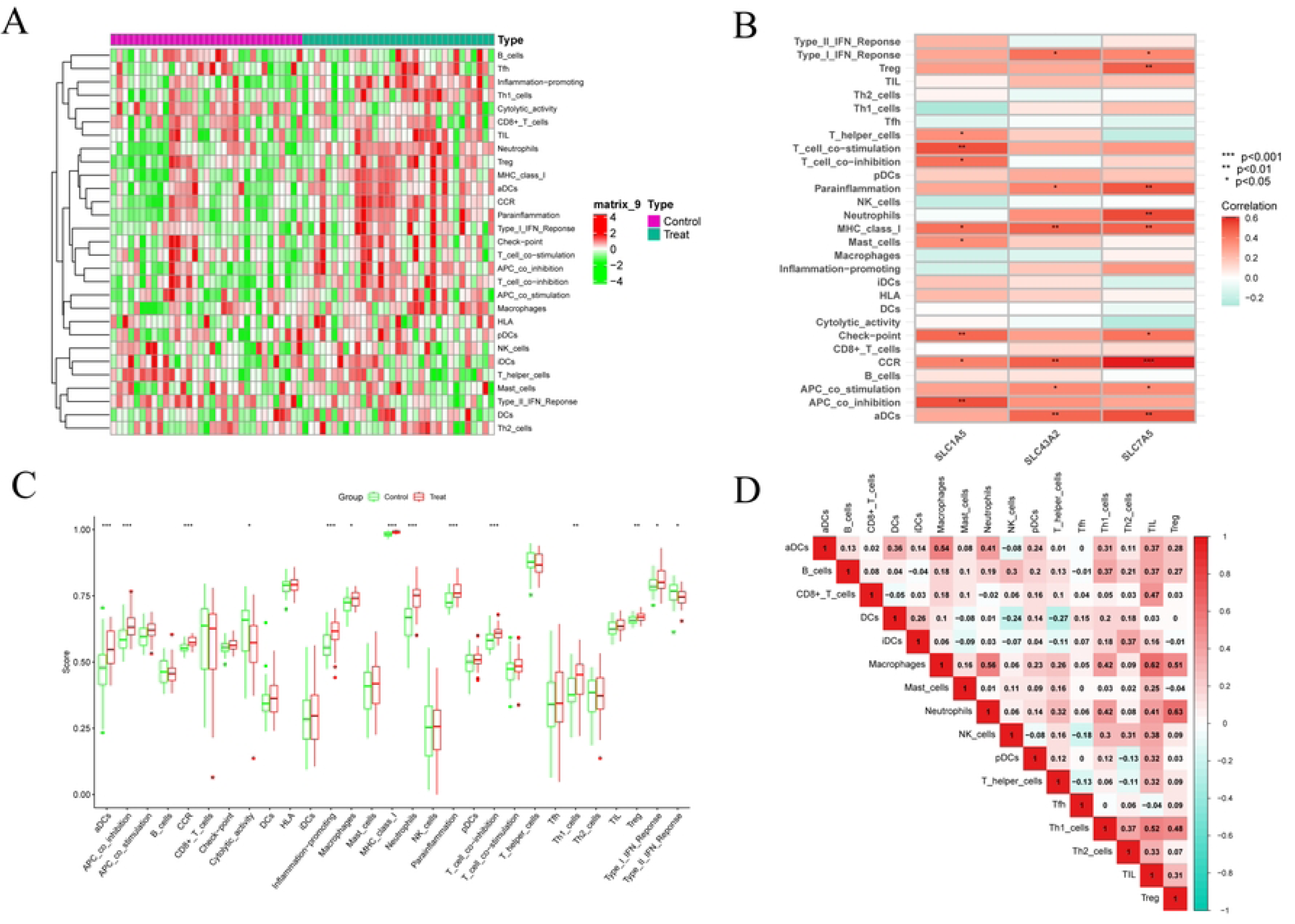
(A) Heatmap of immune cell infiltration between HIRI and control samples, with red and green representing high and low expression, respectively. (B) The correlation between 29 immune cells and functions and three marker genes. Red and green colors represent positive and negative correlations, respectively. (C) Boxplots indicated the differences in immune cells and function between HIRI and control samples. (D) Correlation analysis among 16 immune cells. p values were showed as: *, p < 0.05; **, p < 0.01; ***, p < 0.001.

### CeRNA networks based on marker genes

To further explore the regulatory mechanisms of the 3 hub genes, we constructed a competing endogenous RNA (ceRNA) network using the TargetScan, miRanda, and miRDB databases. The analysis identified 3 mRNAs, 76 miRNAs, and 150 lncRNAs (Figure 8A). Our results indicated that 128 lncRNAs can regulate the expression of SLC43A2 through competitively binding of hsa-miR-149-3p, hsa-miR-1976, hsa-miR-612, hsa-miR-1207-5p and hsa-miR-129-5p. Among them, 19 shared lncRNAs targeted has-miR-149-3p and 17 shared lncRNAs targeted has-miR-1976. A total of 38 lncrnas were able to competitively bind to 7 mirnas, Including hsa-miR-612, hsa-miR-185-3p, hsa-miR-615-5p, hsa-miR-875-3p, hsa-miR-486-3p, hsa-miR-92b-5p, and hsa-miR-194-5p, and subsequently regulate SLC7A5. Among them, 16 lncrnas targeted hsa-miR-612 and 7 lncrnas targeted hsa-miR-185-3p. For SLC1A5, there are two regulated mirnas, including hsa-miR-361-3p and hsa-miR-2355-5p. In addition, both hsa-miR-486-3p and hsa-miR-612 were involved in regulating the expression of SLC43A2 and SLC7A5. Notably, seven miRNAs, including has-miR-541-3p and has-miR-31-5p, could simultaneously bind lncRNA C10orf91 to regulate the expression of SLC7A5 and SLC43A2.

**Figure 8.**
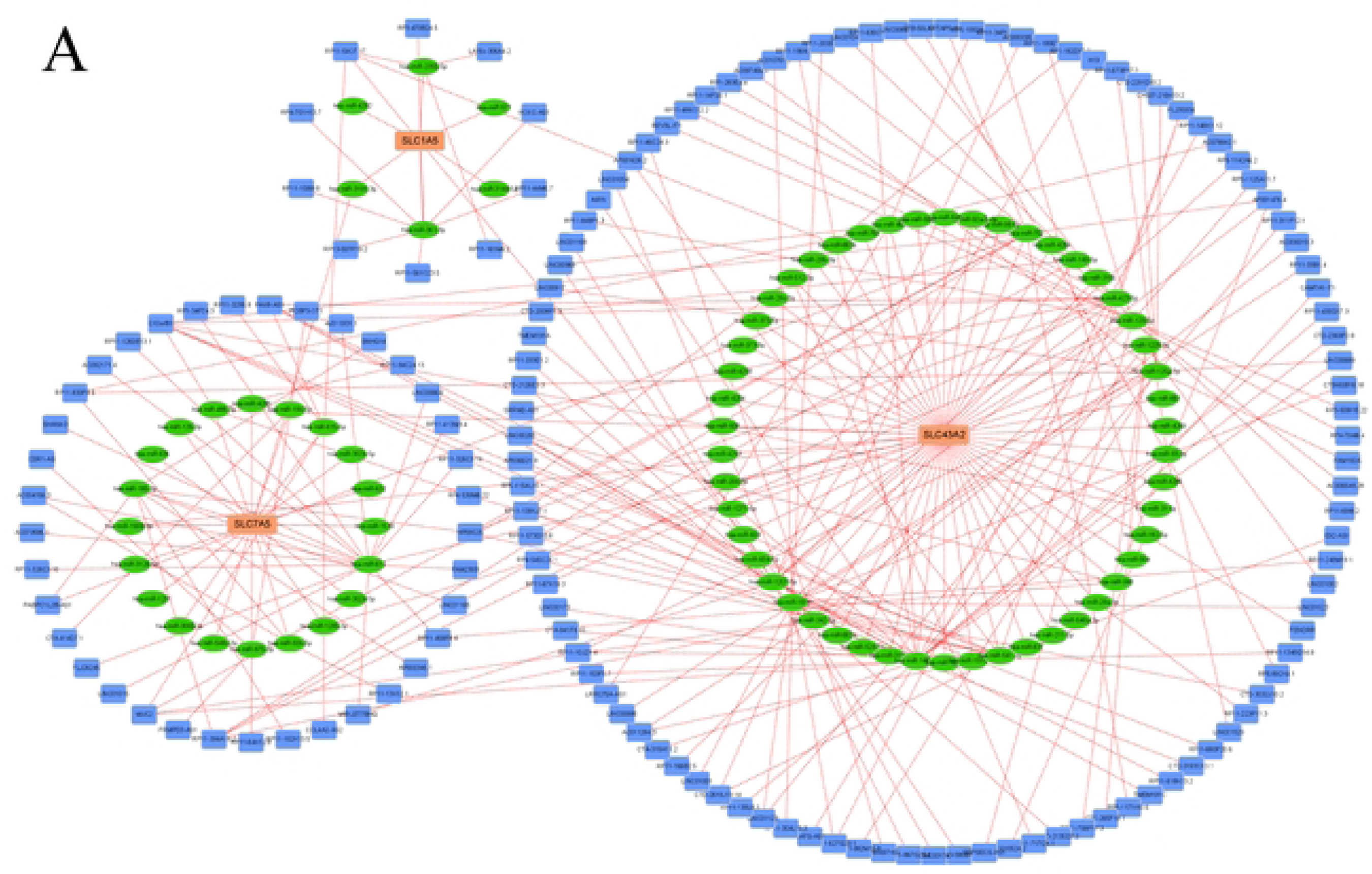
CeRNA networks based on marker genes. (A) The ceRNA network based on marker genes. With Pink dots for mRNA, green dots for miRNA, and blue dots for lncRNA.

### Altered expression of DE-BRGs in HIRI

AST and ALT measurements revealed a significant increase in the HIRI group compared to the control group (Figure 9A-B). Furthermore, HE staining showed pronounced liver damage in the HIRI group, characterized by a marked loss of liver architecture, disintegration of hepatic cords, and red blood cell exudation from the hepatic cords (Figure 9C). Collectively, these results confirm the successful establishment of the HIRI model.

**Figure 9.**
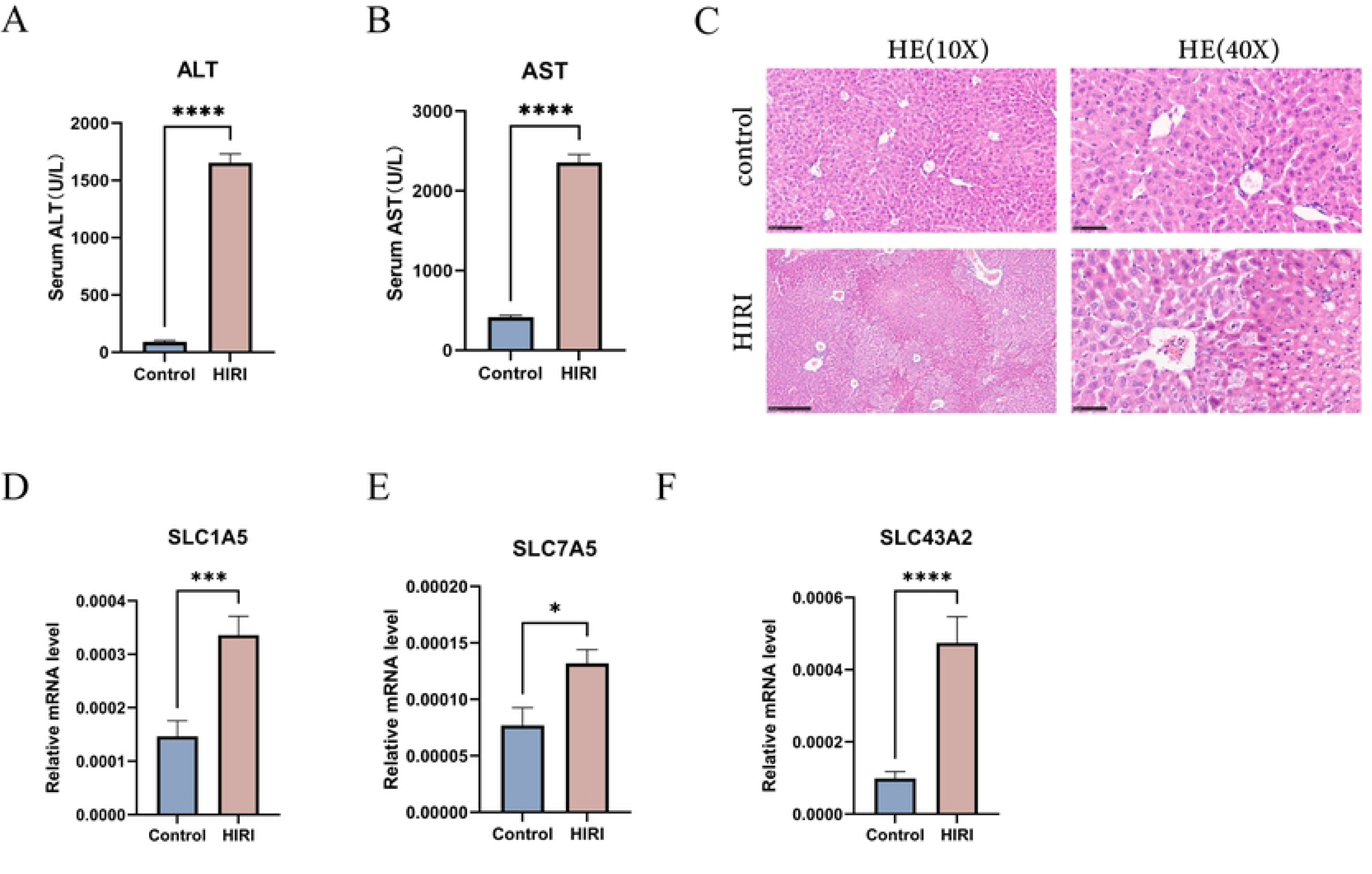
HIRI mice model construction and Altered expression of DE-BRGs in HIRI. (A, B) serum alanine aminotransferase (ALT) and aspartate aminotransferase (AST) value in both wildtype and HIRI mice. (C) The HE staining of wild type and HIRI mice. The result of HE staining showed significant liver injury in HIRI group. (D-F) mRNA expression of SLC1A5, SLC7A5 and SLC43A2 by RT-PCR. Liver tissues and blood of the wildtype and HIRI mice were collected. p values were showed as: *, p < 0.05; **, p < 0.01; ***, p < 0.001. DE-BRGs, differential expression of branched-chain amino acid metabolism-related genes. Wildtype control mice (n=8), HIRI mice(n=8).

To further elucidate the role of BRGs in HIRI, we assessed the mRNA expression levels of SLC7A5, SLC1A5, and SLC43A2 using qRT-PCR. The results demonstrated a significant upregulation of these 3 hub genes in HIRI mice compared to control mice (Figure 9D-F). These findings suggest that BRGs play a pivotal role in HIRI development, thereby supporting their potential as key regulators in HIRI progression.

## Discussion

Ischemia-reperfusion injury is a severe condition that necessitates medical intervention to limit cellular damage and maintain organ function^20^. It frequently arises in medical procedures such as hemorrhagic shock resuscitation, liver transplantation, and partial liver resection, and plays a significant role in liver damage, which can ultimately result in organ failure^21^. From a mechanistic viewpoint, mitochondrial dysfunction is the driving force behind hepatic ischemia-reperfusion injury (HIRI), contributing to oxidative stress, metabolic imbalances, inflammation, and immune dysregulation^22^. BCAAs are vital for mammalian growth, and impaired BCAA metabolism has been recognized as a major risk factor for coronary artery disease, diabetes, heart failure, and myocardial ischemia-reperfusion injury^23–25^. Nonetheless, the involvement of BCAA metabolic pathways in the development of HIRI has not been thoroughly investigated. Consequently, this study aims to uncover BCAAs associated with HIRI and assess the potential diagnostic and therapeutic relevance of DE-BRGs in HIRI.

In this study, we analyzed the differential expression of BRGs in HIRI and control liver samples obtained from the GEO database, ultimately identifying 13 DE-BRGs associated with BCAA metabolism. Compared to healthy individuals, the abnormally BRGs expressed in HIRI patients, suggesting their critical role in the onset and progression of HIRI. Subsequently, we analyzed the correlations among DE-BRGs, revealing that some exhibit significant synergistic or antagonistic interactions, indicating their mutual regulatory roles in HIRI pathogenesis. At the protein level, several pairs, including ECHS1-ABAT and SLC1A5-SLC43A2-SLC7A5-SLC3A2, as well as the AUH-ACAD8-BCKDHB-DLD-IVD-ACADSB-MCCC1-ECHS1 group, were found to be closely related, suggesting extensive interactions between BRGs at both the gene and protein levels during HIRI development.

GO and KEGG enrichment analyses were conducted to explore the potential functions of DE-BRGs in HIRI. The findings primarily indicate associations with amino acid catabolism, the mTOR signaling pathway, and the TCA cycle. This aligns with earlier studies showing that BCAAs stimulate mTOR mRNA translation, regulating protein synthesis at the molecular level.^10^. Meanwhile, BCKDHA, a metabolic enzyme involved in BCAA catabolism, is regulated by the key enzyme BCKDK^26,27^. In α-cell-specific BCKDHA knockout mice, Yang et al. demonstrated that impaired BCAA catabolism could reduce mTOR signaling, thereby inhibiting α-cell proliferation^28^. Zhai et al. similarly noted that APN promotes ERK signal transduction in HCC cells by mediating BCKDK phosphorylation, which enhances HCC proliferation and metastasis^29^. However, our results reveal that the key regulators of branched-chain amino acid metabolism, BCKDHA and BCKDK, did not exhibit significant differences between HIRI and control liver samples, diverging from previous studies^29,30^. Therefore, further research is required to clarify the relationship between DE-BRGs and BCAA metabolism in HIRI. Additionally, GO analysis of the cellular component emphasizes that these 13 DE-BRGs primarily function within the mitochondrial matrix. This aligns with Kitagawa’s study, which highlights the critical role of mitochondria in maintaining cellular homeostasis, particularly in processes like apoptosis, inflammation, and ROS synthesis^31^. Under stress conditions such as hypoxia or cytotoxin activation, mitochondrial membrane integrity is compromised, triggering the pro-apoptotic Bcl-2 family and exacerbating ischemia-reperfusion injury^32,33^. These findings suggest that mitochondrial dysfunction plays a crucial role in HIRI.

Machine learning algorithms, which remove redundant factors and retain only variables relevant to the outcome, have seen growing use in medical research^34–36^. In our study, we applied three machine learning algorithms—LASSO, Random Forest (RF), and SVM-REE—to identify three hub genes (SLC7A5, SLC1A5, SLC43A2) that accurately predict the risk of hepatic ischemia-reperfusion injury (HIRI), achieving an AUC value of 0.834. We further validated the model using an external dataset (GSE15480), where the AUC value reached 1.000. Additionally, in the validation set, the AUC values for SLC7A5, SLC1A5, and SLC43A2 each exceeded 0.85, highlighting their strong predictive value as sensitive biomarkers for HIRI. Using these three hub genes, we constructed a nomogram model, along with a calibration curve and decision curve analysis (DCA), to further validate the diagnostic accuracy and clinical relevance of the model. Although the calibration curve showed close alignment with the standard curve, it did not yet reach an optimal level, possibly due to the limited sample size, which requires further investigation.

Long non-coding RNAs (lncRNAs), functioning as competitive endogenous RNAs (ceRNAs), can bind to miRNAs competitively, thereby regulating mRNA expression and influencing physiological interactions between various cell types^37^. Based on the potential interaction within the lncRNA-miRNA-mRNA axis, we constructed a ceRNA network for HIRI. Our findings revealed that 19 shared lncRNAs target hsa-miR-149-3p to regulate SLC43A2 expression. Ma et al. discovered that microcystin-LR (MC-LR) promotes the expression of hsa-miR-149-3p and may contribute to MC-LR-induced hepatitis and liver cancer through the MAPK pathway^38^. Additionally, 16 shared lncRNAs target hsa-miR-612 to regulate SLC7A5 expression. Lu et al. reported that circETFA and CCL5 competitively bind to hsa-miR-612, promoting hepatocellular carcinoma (HCC) progression by modulating the PI3K/Akt signaling pathway.^39^. These results suggest that lncRNAs may regulate the expression of three marker genes (SLC7A5, SLC1A5, and SLC43A2), offering new insights into the pathogenesis of HIRI. However, the precise mechanisms involving these characteristic genes in HIRI require further validation through in vitro and in vivo studies.

SLC7A5 encodes LAT1, a sodium-independent transporter primarily responsible for transporting BCAAs^40^. SLC7A5 encodes LAT1, a sodium-independent transporter primarily responsible for transporting branched-chain amino acids (BCAAs)^41–43^. Additionally, increased levels of activating transcription factor 4 (ATF4) upregulate SLC7A5, leading to greater uptake of isoleucine and leucine, which in turn activate mTORC1 and inhibit autophagy^44^. In this study, we observed increased mRNA expression of SLC7A5 in HIRI mice compared to controls, although further protein-level validation is needed. GSEA and GSVA indicated that SLC7A5 participates in several immune and inflammatory processes, including cytokine-cytokine receptor interactions, MAPK signaling, NOD-like receptor signaling, and interleukin-21 production. Members of the MAPK family include ASK1, TAK1, ERK1/2, JNK, and p38^45^. Deng et al. demonstrated that BPS reduces inflammation, apoptosis, and autophagy by inhibiting p38 and JNK pathways, thereby alleviating HIRI^46^. Similarly, Hou et al. showed that the MAPK signaling pathway mediates the activation of the NLRP3 inflammasome, triggering pro-inflammatory cytokine secretion and hepatocyte damage^47^. Thus, further investigation into the relationship between SLC7A5 and BCAA metabolism in HIRI is essential for future research.

The SLC1A5 protein functions as a transporter that facilitates the absorption of neutral amino acids and plays a crucial role in glutamine uptake^48^. As an essential energy source for the enterohepatic circulation, elevated intestinal glutamine levels can increase blood concentrations of αKG, which promotes macrophage M2 polarization by enhancing the TCA cycle to repair HIRI. In our study, we observed that SLC1A5 expression was higher in HIRI samples compared to control liver samples, indicating a positive correlation between SLC1A5 expression and HIRI. GSEA and GSVA revealed that SLC1A5 is involved in ECM receptor interaction, mTOR signaling pathways, and N-polysaccharide biosynthesis in pre-ribosomes. Similar to SLC7A5, SLC1A5 regulates mTORC1 signaling activation, leading to increased phosphorylation of mTORC1 and subsequent regulation of cell growth and proliferation^49^. Wang J’s study demonstrated that suberoylanilide hydroxamic acid alleviated orthotopic liver transplantation (OLT)-induced IRI by upregulating autophagy in Kupffer cells via the AKT/mTOR signaling pathway. Additionally, SLC1A5 has been shown to be upregulated in various tumors, including melanoma, lung cancer, colon cancer, and breast cancer, supporting the significantly increased energy demands of tumor cells compared to normal cells^50–53^.

SLC43A2 (LAT4) is a uniporter that transports large, essential neutral amino acids, primarily BCAA^54^. It is highly expressed in tissues such as the placenta, peripheral blood leukocytes, kidneys, small intestine, and brain. Bian et al. found that tumor cells outcompete T cells for methionine uptake through SLC43A2, which affects histone methylation and T cell function. Inhibiting SLC43A2 restores T cell function, thereby enhancing anti-tumor immunity in preclinical models^55^. Peng H et al. discovered that in esophageal squamous cell carcinoma (ESCC), SLC43A2 activates the NF-κB signaling pathway by increasing methionine uptake, creating a positive feedback loop that further upregulates SLC43A2 expression and promotes ESCC progression^56^. GSEA and GSVA revealed that SLC43A2 is involved in several immune processes, including the MAPK, NOD-like receptor, and chemokine signaling pathways. In our study, we observed elevated SLC43A2 expression in liver samples from the HIRI group compared to the control group, consistent with previous reports^57,58^. Additionally, both SLC43A2 and SLC7A5 are associated with the cytokine-cytokine receptor interaction pathway, involving the TGF-β and TNF families^58^, which are critical components of the inflammatory process and play a pivotal role in HIRI. These findings indicate that the three core genes identified play multifaceted roles in the occurrence and progression of HIRI. Understanding the mechanisms of these genes is essential for advancing therapeutic strategies for HIRI.

In the early stages of hepatic ischemia-reperfusion injury (HIRI), dysregulation of liver mitochondrial function and microcirculation leads to hepatocyte injury, triggering a cascade of immune responses that worsen liver damage by recruiting and activating macrophages, neutrophils, T cells, and monocytes^59,60^. Our ssGSEA of immune infiltration in HIRI patients revealed a significant increase in the proportion of macrophages, neutrophils, aDCs, Treg, and Th1 cells compared to controls. These results suggest that the three hub genes—SLC7A5, SLC1A5, and SLC43A2—are associated with various immune cells and functions, particularly neutrophils, aDCs, and Tregs, aligning with prior research^61^. KCs, play a key role in HIRI. In the early ischemic stage, KCs are activated first, secreting cytokines such as TNF-α and IL-1β, which recruit additional immune cells, thereby exacerbating liver injury^62^. Moreover, recent studies show that DAMPs released from necrotic hepatocytes activate inflammasome components such as NLRP3 and AIM2 in KCs via pattern recognition receptors, further intensifying HIRI^63^. Our findings also confirmed a significant increase in macrophage infiltration in the HIRI group compared to controls Both SLC43A2 and SLC7A5 were strongly associated with aDCs. DCs are highly potent professional antigen-presenting cells, and research by Castellaneta et al. demonstrated that severe depletion of plasmacytoid DCs (pDCs) in mice downregulated IFN-α expression in liver tissue, attenuating HIRI^64^. This highlights the damage-inducing and pro-inflammatory role of DCs.. However, Nakamoto et al. found that loss of the prostaglandin E receptor EP3 (PTGER3) in conventional DCs (cDCs) increased macrophage activity and delayed liver repair after warm ischemia, indicating that the roles of DCs in HIRI may vary depending on their subtypes^65^. This variability suggests a direction for future research.. Neutrophils also play a significant role in liver injury. Kaltenmeier et al. showed that DAMPs activate neutrophils, leading to the release of enzymes that activate the complement system and trigger an inflammatory response during HIRI^66^. Neutrophils also play a significant role in liver injury. Kaltenmeier et al. showed that DAMPs activate neutrophils, leading to the release of enzymes that activate the complement system and trigger an inflammatory response during HIRII^67^. Targeting neutrophils may represent a promising therapeutic approach to alleviating HIRI. CD4+ T cells are critical mediators of inflammation in the HIRI cascade, capable of activating macrophages and amplifying the immune response^68,69^. CD4+ T cells can differentiate into various subtypes, including Th1, Th2, Th17, and Treg cells^70^. Th1 cells primarily eliminate intracellular pathogens by secreting IFN-γ and recruiting immune cells, while Treg cells are involved in immunosuppression through the secretion of IL-10 and TGF-β. Our study showed that SLC7A5 is significantly associated with Treg cells. In summary, these findings underscore the crucial role of the immune system in the development and progression of HIRI.

We also constructed a prediction model for HIRI and explored the characteristic genes and signaling pathways involved. However, our study has several limitations. First, the data were obtained from the GEO public database, which restricts sample size and introduces potential selection bias due to the absence of raw sequencing data. Second, although our mice experiments partially validated the bioinformatics analysis, we were unable to collect sufficient clinical samples due to time constraints. Gathering adequate clinical samples is vital to fully uncover the role of BCAA metabolism in HIRI. Additionally, the discrepancy between RNA sequencing and qRT-PCR results highlights the complexity of BCAA metabolic regulation in HIRI. Therefore, future research should aim to comprehensively investigate these mechanisms to better understand the regulatory pathways involved.

## Conclusion

In this study, we identified three characteristic genes through bioinformatics analysis and machine learning algorithms that may serve as potential diagnostic biomarkers for HIRI. Additionally, we demonstrated the critical role of the immune system in the occurrence and progression of HIRI, emphasizing significant differences in immune responses between HIRI and control liver samples. Our findings provide new insights into the role of branched-chain amino acid metabolism in HIRI and establish a foundation for future research into the pathogenesis of HIRI and potential therapeutic targets.

## Acknowledgments

The authors sincerely express gratitude for the insightful comments provided by the reviewers on the present article. Additionally, we would like to thank the researchers who generously contributed the datasets (GSE23649 and GSE15480).

## Author contributions

The formal analysis and original draft of the manuscript were performed by JZ, MW. Project administration was performed by SJL, SL, ML, QL. Software analysis was performed by JZ, MW, GO. Data curation was conducted by JZ, SJL, GP, HX. The experiment was performed by MW, ML, QL. The writing, reviewing, and editing of the article were contributed by JZ, GO, GP, HX. All authors contributed to the article and approved the final version for submission.

## Ethics statement

All animal experiments were approved by the Animal Care and Use Committee of Liuzhou People’s Hospital Affiliated to Guangxi Medical University (IACUC20240028). All experiments were performed in accordance with regulations and complied with the ARRIVE guidelines.

## Data availability statement

The datasets utilized in this study were obtained from the GEO database (https://www.ncbi.nlm.nih.gov/geo/) with the following data accessions: GSE23649 and GSE15480. The original codes used for analyses presented in the study are publicly available. This data can be found here: https://github.com/jiahui980702/HIRI-BCAA-.

## Abbreviations

AUC: area under the curve
CC: cellular component
DEGs: Differentially expressed genes
DCA: decision curve analysis
GEO: Gene Expression Omnibus
GO: Gene Ontology
GSEA: gene set enrichment analysis
GSVA: gene set variation analysis
HIRI: Hepatic ischemia and reperfusion injury
KEGG: Kyoto Encyclopedia of Genes and Genomes
LASSO: least absolute shrinkage and selection operator
PPI: protein protein interaction
RF: random forest
RFE: support vector machine recursive feature elimination
ROC: receiver operating characteristics
SVM-ssGSEA: single-sample gene set enrichment analysis
TCA: tricarboxylic acid

